# A spatial-temporal map of glutamatergic neurogenesis in embryonic cerebellar nuclei uncovers a high degree of cellular heterogeneity

**DOI:** 10.1101/2023.10.22.563467

**Authors:** Filippo Casoni, Laura Croci, Francesca Marroni, Giulia Demenego, Ottavio Cremona, Franca Codazzi, G. Giacomo Consalez

## Abstract

The nuclei are the main output structures of the cerebellum. Each and every cerebellar cortical computation reaches several areas of the brain by means of CN processing and integration. Nevertheless our knowledge of these structures is still limited compared to the cerebellar cortex. Here, we present a genetic inducible fate mapping study characterizing rhombic lip-derived glutamatergic neurons of the nuclei, the most conspicuous family of long-range cerebellar efferent neurons. Glutamatergic neurons mainly occupy dorsal and lateral territories of the lateral and interposed nuclei, as well as the entire medial nucleus. They are born starting from about embryonic day 9.5, with a peak between 10.5 and 12.5, and invade the nuclei with a lateral to medial progression. While some markers label a heterogeneous population of neurons sharing a common location (Brn2), others appear to be lineage specific (Tbr1, Lmx1a, Meis2). A comparative analysis of Tbr1 and Lmx1a distributions reveals an incomplete overlap in their expression domains, in keeping with the existence of separate efferent subpopulations. Finally, some tagged glutamatergic progenitors are not labeled by any of the markers used in this study, disclosing further complexity. Taken together, our results obtained in late embryonic nuclei shed light on the heterogeneity of the excitatory neuron pool, underlying the diversity in connectivity and functions of this largely unexplored cerebellar territory. Our findings lay the groundwork for focused functional analyses of individual subpopulations of nuclear neurons.

## Introduction

The function of the cerebellum requires an interplay between its cortex and output circuitry, represented almost entirely by the CN. A broad range of inputs from the entire central nervous system is elaborated by the cerebellar cortex ^1^ and forwarded to the CN, which project their output mainly to the brainstem and diencephalon.

In both mice and humans, the CN consist of a heterogeneous collection of neurons positioned deep within the cerebellar hemispheres. Although there are differences in size and complexity between the mouse and human CN, the general organization and function are well-conserved across mammalian species. The human CN comprise four separate nuclei named fastigial, globose, emboliform and dentate, from medial to lateral ^1,2^. Instead, rodent cerebella contain three separate nuclei, since the globose and emboliform nuclei are anatomically fused, giving rise to the so-called interposed nucleus. Thus, from medial to lateral, the rodent CN consist of the medial (MED), interposed (INT) and lateral (LAT) ^3,4^.

Anatomically, it is now evident that the individual nuclei can be further subdivided into various subnuclei, which might reflect the modular organization of the olivo-cortico-nuclear system ^5,6^. Each of these subnuclei plays its own role and has its set of connections, although it remains largely unknown how these subnuclei exert their function, as the classification of neuronal types making up the CN subnuclei is still in progress. Two or three classes of CN neurons were historically described merely based on their soma sizes ^3,7,8^ and the synthesis and release of glutamate or GABA ^9-^ 14_._

Recently, CN neurons were subdivided in five classes reviewed in ^1^. This classification was based on different lines of evidence including neurotransmitter production, morphology, location, and transcriptomic signatures identified in adult mice. According to this subdivision, the CN contain two types of glutamatergic projection neurons, referred to as class A and class B, which exhibit different morphological features and transcriptomic signatures ^1,15^. Additionally, three classes of GABAergic or GABA-and glycinergic neurons were identified, differing with respect to physiology and developmental origin ^9,15,16^.

A robust link between CN neuron progenitors and their mature counterparts is still missing or is largely incomplete. Significant progress has been made in understanding the development of glutamatergic neurons populating the CN, through the use of *Atonal basic helix-loop-helix Transcription Factor 1* (*Atoh1*)-tagged mouse cerebella ^17,18^. Class-A and Class-B glutamatergic projection neurons of the CN derive from *Atoh1*+ rhombic lip (RL) progenitors mostly between E10.5 and E12.5. They migrate as part of the so-called rostral RL migratory stream, tangentially along the surface of the cerebellar primordium ^17^, and reach the nuclear transitory zone (NTZ) as early as E11.5 ^17,18^. While migrating, neurons downregulate *Atoh1* and upregulate several transcription factors, including Pax6 (E13.5) ^19^ and Pou3f1 (E10.5) ^20^. In the NTZ, the nascent CN express Tbr2 and Tbr1 ^19^, which mark a subset of neurons of the MED, as well as Brn2 (Pou3f2), Olig2 and/or Irx3, which are thought to mainly give rise to LAT and INT neuronal subsets ^20^.

The present genetic inducible fate mapping (GIFM) study expands the analysis of progenitors derived from the *Atoh1* lineage and describes the distribution of molecularly defined derivatives of this lineage in the perinatal CN. Our GIFM results and the availability of the Atoh1-CreER^T2^ line will allow the dissection of developmental trajectories that link embryonic and adult lineages in the CN

## Results

To investigate the cellular composition of the mouse CN, we utilized a tamoxifen (Tam)-inducible *Atoh1::CreER^T^*^2^*/0* transgenic mouse line ^18^ mated with a robust Cre-inducible *tdTomato* (tdT)-reporter line ^21^. *Atoh1* is expressed in the rhombic lip and is essential for the development of glutamatergic neurons of the cerebellum ^17,18^.

GIFM allows temporal control of Cre-mediated recombination for a ∼24 hr time window subsequent to Tam injection. Tam (70mg/kg) was administered via oral gavage to pregnant dams at E9.5, E10.5, E11.5, E12.5, or E13.5. In this paper, cerebella were analyzed at E18.5 and P60.

### Distribution of glutamatergic lineage neurons in the late embryonic CN

Upon Tam administration at E9.5 and immunofluorescence analysis at E18.5 (hereon TamE9.5), we only observed tdT fluorescence in a small number of cells restricted to the dorsal area of the presumptive LAT and INT CN (see red dotted lines), in rostral sections (**Fig.1A**), while hardly any cells were tagged in caudal sections (**Fig.1B**).

**Fig.1.**
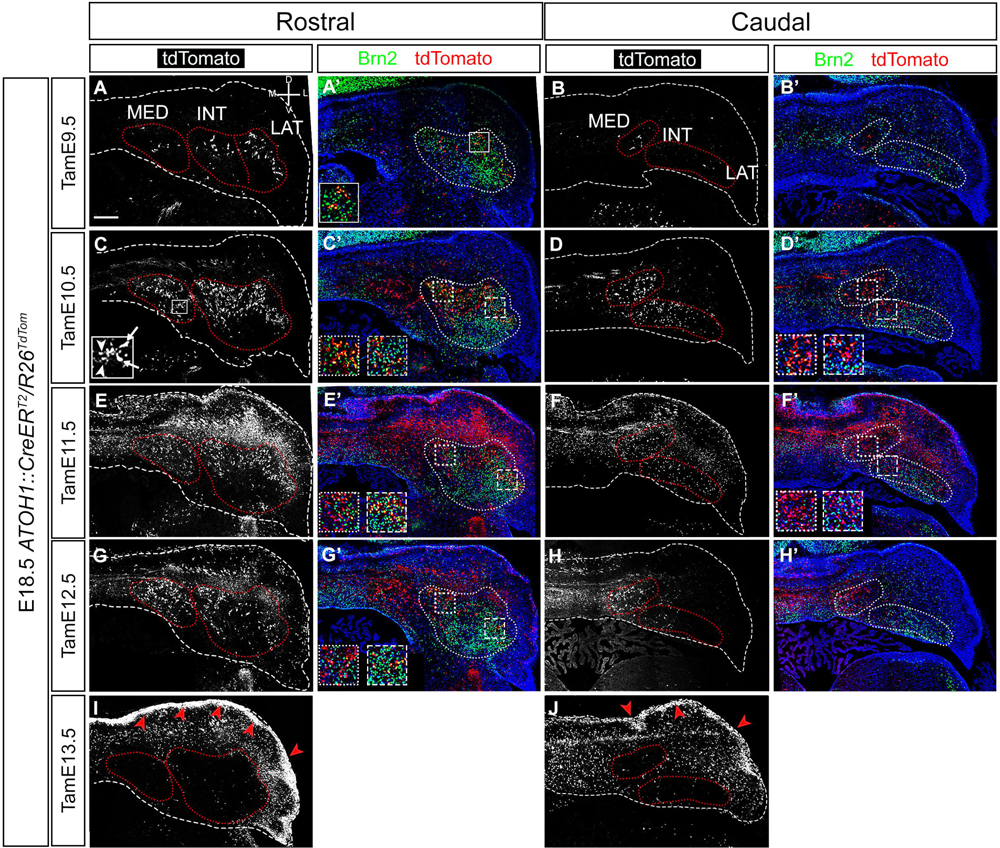
*Atoh1 lineage progenitors migrate into the embryonic CN from TamE9.5 to TamE12.5 in a lateral to medial progression.* The perimeters of cerebella and of each CN are delimited by dashed and dotted lines, respectively. **A,C,E,G,I.** Anti-tdT immunofluorescence in rostral hemisections of E18.5 cerebella; *Atoh1* lineage neurons are visible in the CN from TamE9.5 to TamE12.5. Only a small number of TamE9.5 neurons of the early-born cohort are detected (**A**) and mostly occupy the dorsolateral area of the LAT. TamE10.5 (**C**) and TamE11.5 (**E**) tdT+ neurons occupy the dorsolateral portion of the LAT, the dorsal area of the INT and the MED. Solid line box in C shows small and large glutamatergic neurons (arrowheads and arrows, respectively). TamE12.5 tdT+ neurons (**G**) are visible in the medial area of the INT and in the entire MED. TamE13.5 tdT+ neurons (**I**) are not visible in the CN. Solid red arrowheads in **I** point to granule cell progenitors in the external granular layer. **A’,C’,E’,G’.** Anti-tdT and Brn2 double immunofluorescence in rostral hemisections of E18.5 cerebella. Brn2+ cells formed compact cellular clusters throughout the LAT and INT. Sparse TamE9.5 tdT+ neurons are also positive for Brn2 in the LAT (solid line box in **A’**) and INT (**A’**). Most of the TamE10.5 tdT+ neurons are Brn2+ in the dorsolateral area of the LAT (dashed box, **C’**) and in the dorsal area of the INT (dotted box, **C’**). TamE11.5 and TamE12.5 tdT+ neurons are also Brn2+ in the lateral portion of the LAT (dashed boxes **E’,G’**) and dorsomedial portion of the INT (dotted boxes **E’, G’**). **B,D,F,H,J.** Anti-tdT immunofluorescence in caudal hemisections of E18.5 cerebella. *Atoh1* lineage neurons are visible in the CN (dotted red lines) from TamE9.5 to TamE12.5. Only sparse neurons occupy the CN in the caudal region of the TamE9.5 cerebellum (**B**). TamE10.5 (**D**) and TamE11.5 (**F**) tdT+ neurons are visible in the MED, INT and LAT. TamE12.5 (**H**) tdT+ neurons are mostly present in the medial area of the INT and in the MED. Virtually no TamE13.5 (**J**) tdT+ neurons are present in the CN. Solid red arrowheads in **J** point to granule cell progenitors in the external granular layer. **B’,D’,F’,H’.** Anti-tdT and Brn2 double immunofluorescence in caudal hemisections of E18.5 cerebella. Only sparse cells were double positive in TamE10.5 in the MED and INT (dotted and dashed boxes, respectively). Virtually all tdT+ cells tagged at E11.5 were negative for Brn2 (boxes in **F’**). In caudal sections, a small number of double positive cells were found in the dorsal portion of the INT (dashed square box) and in the MED (dotted square box) (**D’, F’**). Size bar in B’: 200 µm.

In rostral sections, Tam administration at E10.5 and E11.5 reveals robust tagging in the dorsal and lateral area of the LAT, in the dorsal and medial area of the INT, and throughout the MED (**Fig.1C, E**). In caudal sections we observed tdT+ cells in the medial portion of the INT after TamE10.5 and throughout the INT after TamE11.5; moreover, diffused signal is observed in the caudal MED (**Fig.1D, F**).

In TamE12.5 rostral sections, tdT+ cell numbers dropped in the dorsal portion of the LAT compared to previous stages of Tam induction: in fact, only few tdT+ cells were tagged in the lateral portion of the LAT (**Fig. 1G**). Robust tdT tagging was observed in the dorsomedial portion of the INT, and throughout the MED (**Fig. 1G**). Tam administration at E12.5 also revealed very few tagged neurons in the caudal INT, while numerous cells were tdT+ in the caudal MED (**Fig1H**).

Finally, in TamE13.5 cerebella, hardly any cells were tagged in rostral or caudal sections of the CN, (**Fig.1I, J**). As expected, numerous cells populating the external granular layer were positive for tdTomato at this stage (**Fig.1I, J** red arrowhead).

At all developmental stages examined, we observed a mixed population of Atoh1-tagged glutamatergic neurons, containing both large (white arrows in Fig. 1 C box) and small cells (white arrowheads in Fig. 1C box), in keeping with observations made in the adult CN ^1,15,16^.

Our data indicate that *Atoh1*-tagged glutamatergic neurons occupy the dorso-lateral and dorsomedial areas of the LAT and INT, respectively at rostral levels of the cerebellar nuclei. The MED, occupied by later-born neurons, display Atoh1-tagged glutamatergic neurons both in rostral and caudal sections. In all nuclei, at E18.5, ventral territories are occupied predominantly by non-glutamatergic neurons (Casoni et al., in preparation).

We further investigated the distribution of the *Atoh1*-tagged glutamatergic neurons in adult rostral sections. Rostrally, the LAT displayed more cells in the TamE10.5 section (**Fig. 2A,A’** dashed cyan lines) compared to TamE11.5 and TamE12.5 sections (**Fig. 2B-C’** dashed cyan lines). Caudally, fewer cells are visible in the MED of TamE10.5 and TamE11.5 sections (**Fig. 2D-E’** dashed cyan lines) compared to TamE12.5 sections (**Fig. 2F,F’** dotted cyan line).

**Fig. 2.**
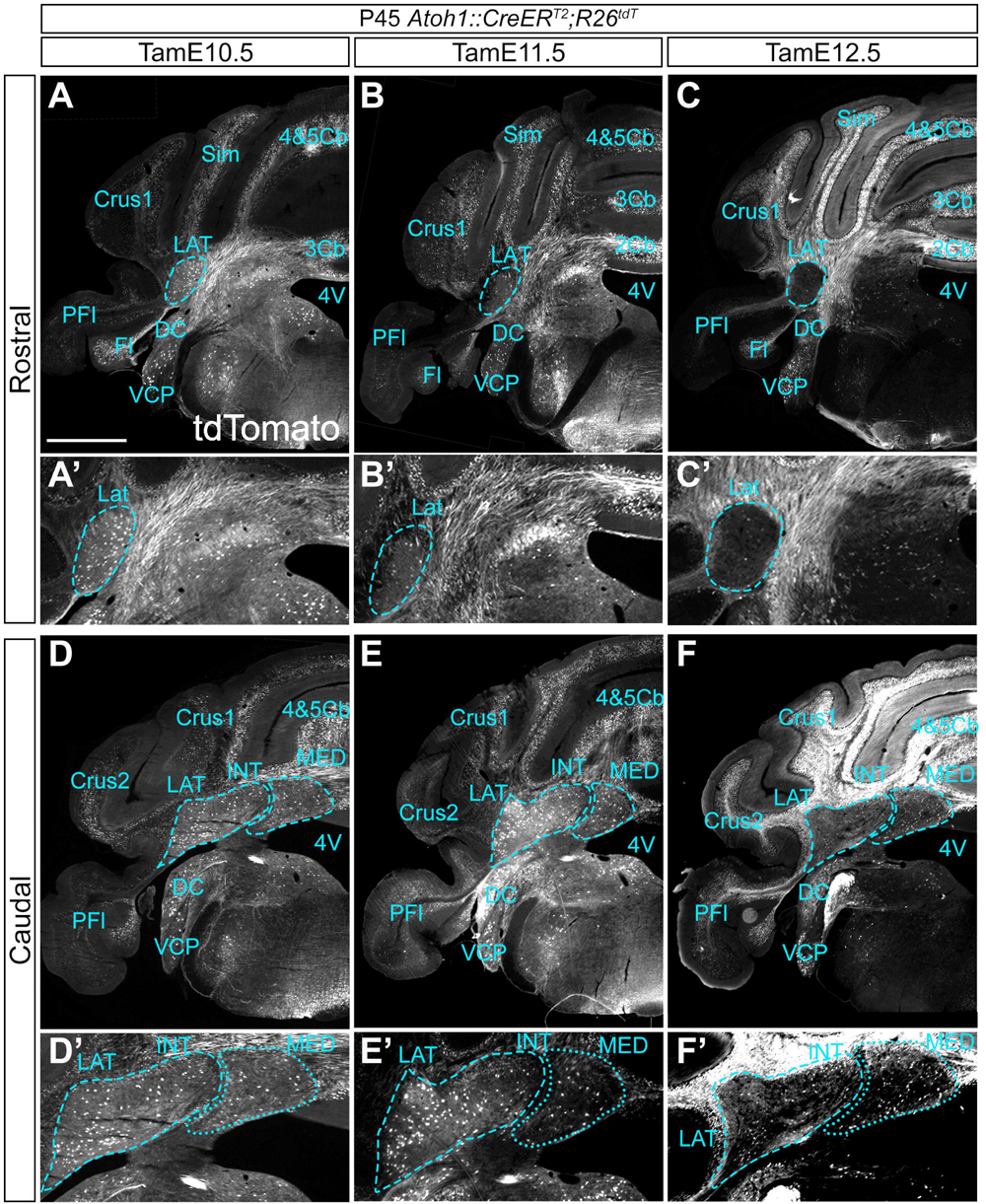
The distribution of *Atoh1* lineage cells in the adult reflects the lateral to medial progression observed in late embryonic sections. The perimeter of each CN is delimited by dashed lines. Immunostaining against tdT in Atoh1::CreER^T2^/R26^tdT^ adult (P60) rostral cerebellar hemisections shows recombination in the LAT (cyan dashed lines); **A-C** (magnification in **A’-C’**). In rostral sections anti-tdT staining shows *Atoh1* lineage neurons in the LAT (cyan dashed lines) from TamE10.5 to TamE12.5. The number of tdT-tagged cells in the LAT decreases from TamE10.5 (**A,A’**) to TamE12.5 (**C,C’**). In caudal sections (**D-F,** magnification in **D’-F’**), anti-tdT staining shows *Atoh1* lineage neurons in the LAT, INT and MED (cyan dashed lines). From TamE10.5 **(D, D’)** to TamE12.5 **(F,F’)**, the number of cells decreases in the LAT and progressively increases in the MED. Crus1: crus 1 of the ansiform lobule, Crus2: crus 2 of the ansiform lobule, DC: dorsal cochlear nucleus, FI: flocculus, Int: interposed nucleus, Lat: lateral nucleus, Med: medial nucleus, PFI: paraflocculus, Sim: simple lobule, VCP: ventral cochlear posterior nucleus, 3Cb: cerebellar lobule III, 4&5Cb: cerebellar lobules IV and V. Size bar: 1mm

### Brn2 labels a heterogeneous population of LAT and INT neurons

To further characterize CN development, we employed established markers of different CN components. Antibodies for Brn2, a POU-domain transcription factor ^22^, label the LAT and INT (not the MED) during late gestation embryogenesis ^19^. We performed dual immunofluorescence for Brn2 and tdT at E18.5 on cerebellar sections from litters treated with Tam at different embryonic stages.

Brn2+ cells form compact cellular aggregates throughout the LAT and INT of both rostral and caudal cerebellar sections. In TamE9.5 CN, very few neurons are double positive for Brn2 and tdT in the LAT and INT (**Fig 1A’,B’**). Instead, Tam administration at E10.5 reveals numerous tdT+/Brn2+ cells in rostral E18.5 sections (**Fig 1C’**), occupying dorsal locations in the INT and LAT (dotted and dashed boxes, respectively). In caudal sections, a tiny number of double positive cells were found in the dorsal portion of the INT and in the MED (**Fig. 1D’**, dashed and dotted boxes, respectively).

Double positive cells were found in the lateral portion of the LAT and dorsomedial portion of the INT in TamE11.5 and TamE12.5 rostral sections (**Fig.1E’,G’**, dashed and dotted boxes, respectively). Hardly any colocalization was found in TamE11.5 and TamE12.5 caudal sections. Interestingly, many tdT+ cells tagged at E11.5 were negative for Brn2, indicating that this protein is not an exhaustive marker of glutamatergic CN neurons (**Fig. 1F’**, both boxes). Finally, no double positive cells were found in E18.5-TamE13.5, either at rostral or at caudal locations (not shown).

### Tbr1 and Lmx1a mark two partially overlapping subpopulations of glutamatergic neurons of the MED

We performed Tbr1 and tdT immunostaining on E18.5 cerebellar rostral sections. Irrespective of the stage of Tam induction, we only observed double labeling in the MED, while Tbr1 is not expressed in LAT and INT at the same stage (**Fig. 3**). Adult medial nuclei comprise a rostral part, a caudal part, and a dorsolateral protuberance (DLP) ^3,23,24^. Accordingly, in the E18.5 MED we identified a ventrolateral mass, a medial mass and a DLP, as sketched in **Fig. 3E**. The DLP was visible only in caudal sections, as described in the adult reviewed in ^1^. Very few Tbr1, tdT double positive cells were found in TamE10.5 rostral and caudal sections (**Fig. 3A,A’**). However, most of the Tbr1, tdT double positive cells were found in rostral sections from TamE11.5 and TamE12.5 cerebella, especially in the ventrolateral portion of the MED, while in the dorsomedial portion of the MED most of the tdT+ cell bodies were Tbr1– (**Fig. 3B,C**). In caudal sections, double positive cells were found in the ventrolateral and medial portion of the MED in TamE11.5 cerebella, and only in the ventrolateral territory of the MED in TamE12.5 sections. No Tbr1+ cells were found in the DLP (**Fig. 3B’,C’)**. Once again, no tdT+ neurons were found in the MED in TamE13.5 CN (**Fig. 3D,D’)**, as described above (**Fig. 1I, J**). Taken together, these data provide significant new details demonstrating that most of the Tbr1+ cells detectable at E18.5 are born between E11.5 and E12.5 and occupy the ventral and ventro-medial position of the MED (see sketch in **Fig. 3E**). We also performed Lmx1a and tdT double immunostaining on E18.5 cerebellar rostral sections. We observed several Lmx1a+ cells in different regions of the cerebellum, including the vermis (presumptive UBCs and granules of lobule X). In the nuclei, most of the Lmx1a, tdT double positive (glutamatergic) neurons were found in the MED (**Fig. 4**), at both rostral (**A-C**) and caudal (**A’-C’**) levels. Lmx1a, tdT double positive cells were found in the MED after Tam induction in a time window spanning E10.5 to E12.5, both in rostral and caudal sections. Anteriorly, most of the double positive cells were tagged at E12.5 in the medial portion of the MED. In caudal sections, double positive cells for Lmx1a and tdT were found throughout the MED, including the DLP, unlike Tbr1+ cells that were excluded from the DLP. Tam administration at E13.5 labeled no tagged neurons in the MED, as previously described (data not shown). These data indicate that at E18.5 most of the Lmx1a+ cells in the CN were tagged between E10.5 and E12.5, and that they occupy a broader area of the nucleus compared to Tbr1+ cells. This, tentatively, identifies two glutamatergic subpopulations of the MED: Lmx1a+, Tbr1+ and Lmx1a+, Tbr1–. To investigate this possibility, we immunostained E18.5 wt rostral sections, using Lmx1a and Tbr1 antibodies (**Fig. 5**). Double immunostaining confirmed previous observations indicating that Tbr1 is located in the medial and ventrolateral area while Lmx1a+ neurons were preferentially distributed in the medial and DLP area (**Fig. 5**). These results confirm the existence of an heterogeneous population of glutamatergic MED neurons inter alia ^23^.

**Fig. 3.**
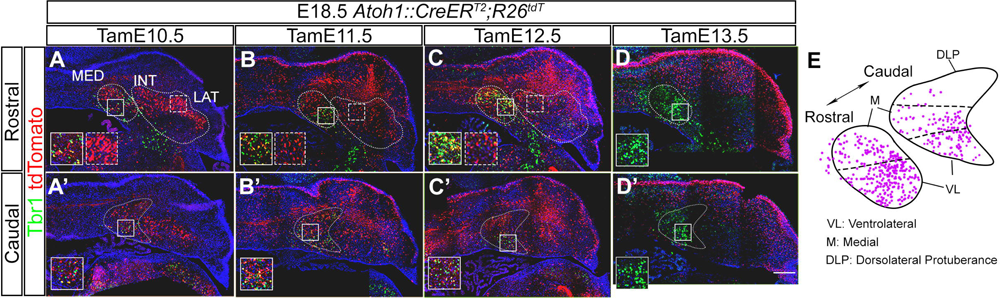
Tbr1+ glutamatergic neurons are localized in the medial and ventrolateral portion of the MED. The perimeter of each CN is delimited by dotted lines. **A-D’.** All sections (E18.5) were immunostained for tdT (red) and Tbr1 (green) **A-D.** In rostral sections, Tbr1+ cells are located in the ventrolateral and medial area of the MED (solid line boxes). No Tbr1 cells can be found in the INT or LAT (dashed line boxes). **A’-D’.** In caudal sections, Tbr1+ neurons are localized in the ventrolateral and medial area of the MED (solid line boxes). **E.** Graphical representation of rostral and caudal sections of the MED: individual Tbr1+ neurons shown in **A-D’** are mapped. The map confirms the presence of Tbr1+ cells in the ventrolateral and medial area of the LAT, and their absence in the dorsolateral protuberance. Size bar: 200 µm

**Fig. 4.**
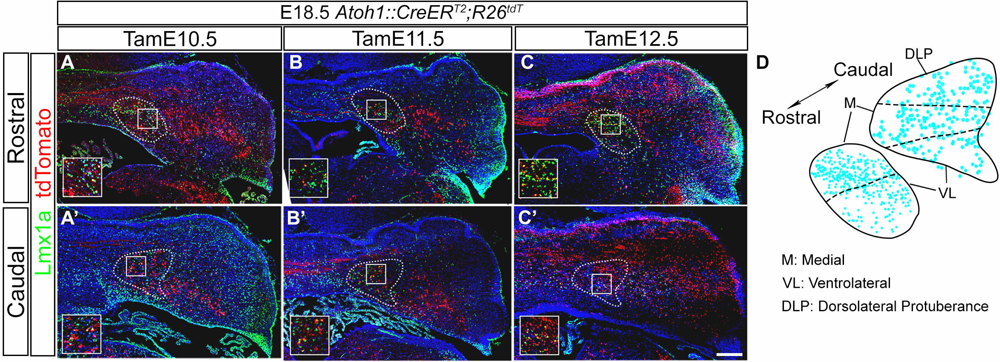
Lmx1a+ glutamatergic neurons are found in the medial area and dorsolateral protuberance of the MED. **A-C’**. The perimeter of the MED is delimited by dotted lines. All sections (E18.5) were immunostained for tdT (red) and Lmx1a (green). **A-C.** Rostral sections: Lmx1a+ cells localize preferentially in the medial area of the MED (solid line boxes). **A’-C’**. Caudal sections: Lmx1a immunostaining is visible throughout the MED, including the DLP (solid line boxes). **D**. Graphical representation of rostral and caudal sections of the MED. Individual neurons that were immunostained for Lmx1a in A-C’ were mapped by superimposing the drawings of the MED to the pictures in A-C’. The map confirms that Lmx1a-positive cells localize mostly in the medial area and in the dorsolateral protuberance of the MED. **A-B’**. Several Lmx1a+ unipolar brush cells are found outside the CN (white arrowheads). Size bar: 200 µm

**Fig. 5.**
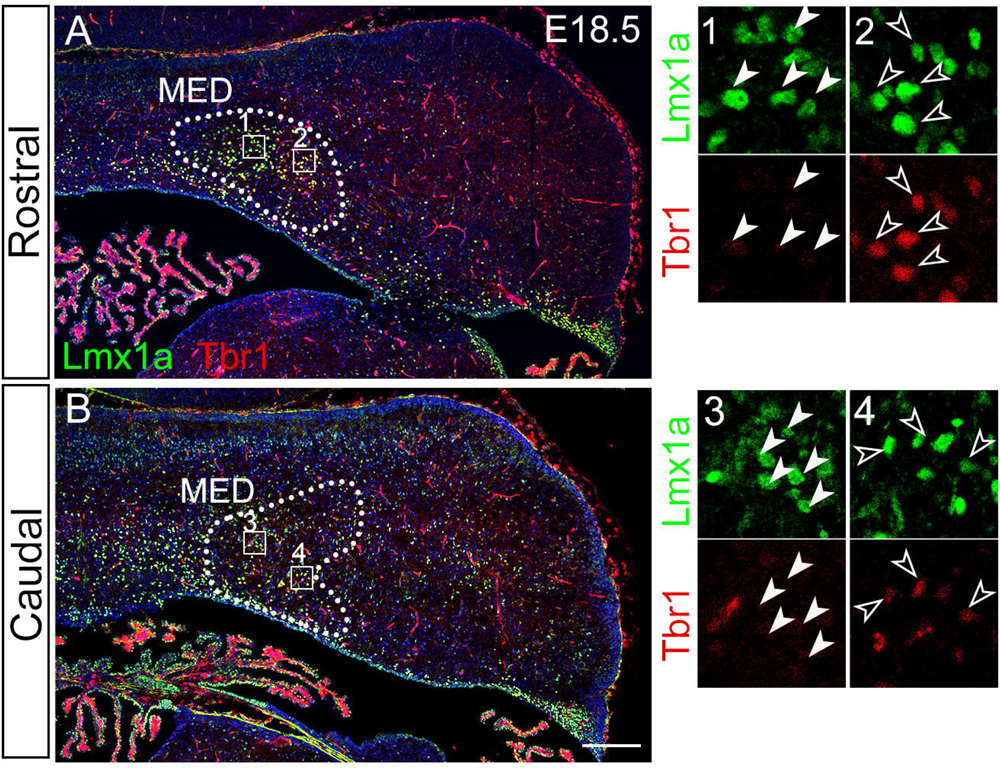
Lmx1a+ and Tbr1+ cell populations are only partially overlapping. **A, B.** Lmx1a and Tbr1 immunostaining in rostral (**A**) and caudal (**B**) sections of the E18.5 MED reveals double positive neurons, as well as cells that are only positive for either marker. Lmx1a+/Tbr1– cells are displayed in boxes 1 and 3 (white arrowheads), while double positive Lmx1a and Tbr1 double positive cells are visible in boxes 2 and 4 (empty arrowheads). Size bar: 200 µm

### Meis2 is a non-exhaustive marker of prenatal glutamatergic CN neurons

*Meis2* belongs to a family of Pbx-related homeobox genes ^25^ and is distributed in expression domains found in all three CN ^26,27^. Again, Meis2 expression in the CN was analyzed by immunostaining (**Fig. 6**) in a series of four E18.5 sections along the rostro-caudal axis. Anteriorly, Meis2+ cells were detected in the dorsolateral area of the LAT and in the dorsal area of the INT **(Fig. 6, A,** dotted lines), while hardly any Meis2+ cells were found in the MED **(Fig. 6, A,** dashed lines). In more caudal sections, however, the expression of Meis2 increased progressively in the medial area of the MED (**Fig. 6, B, C**, dashed lines) and reached its highest levels in caudal most sections, at the level of the DLP **(Fig. 6, D,** dashed line). Using Abs specific for Meis2 and tdT at different stages of Tam induction, we showed that most of the *Atoh1* lineage cells were positive for Meis2 in the dorsal and, especially, in the lateral portion of the LAT, in addition to the dorsomedial portion of the INT in rostral sections (**Fig. 7** **A-C)**. Our data indicate that the bulk of Meis2+ glutamatergic neurons are born at E11.5 (**Fig. 7** **B,B’)**. Meis2 protein distribution at E18.5 suggests that this marker labels a specific subpopulation of glutamatergic neurons, since numerous neurons belonging to the glutamatergic lineage are Meis2-negative at this stage.

**Fig. 6.**
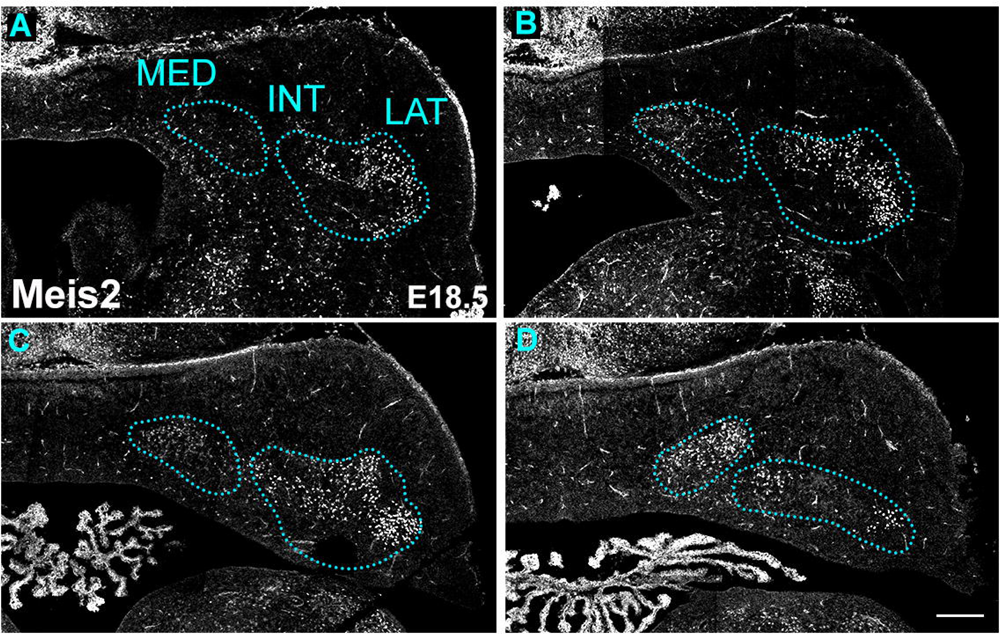
In the MED Meis2 expression is restricted to caudal territories. Meis2 immunostaining of a rostro-caudal series of frontal sections (from **A** to **D**). In **A** Meis2 is expressed in the LAT and in the dorsomedial area of the INT, and it is excluded from the MED. More caudally, in **B**, Meis2 expression is detected more ventrally in the LAT and in the dorso-lateral area of the INT; only a weak positivity is visible in the MED. **C**. Meis 2 expression is visible dorso-and ventrolaterally in the LAT, medially in the INT and weakly in the MED. Finally, in **D** Meis2 is strongly expressed in the DLP of the MED and in a small number of cells of the INT and LAT. Size bar: 200 µm

**FIG. 7.**
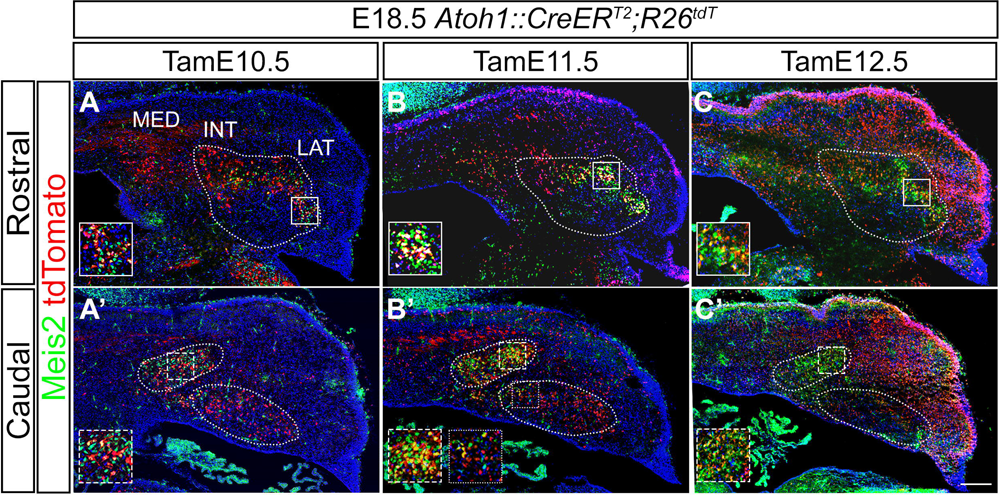
The majority of Meis2+ glutamatergic neurons are born at E11.5. **A-C’**. All sections (E18.5) were immunostained for tdT (red) and Meis2 (green). Very few Meis2, tdT double positive cells are visible in TamE10.5 (insets in **A,A’**). The bulk of Meis2+ neurons are tagged at E11.5 (insets in **B,B’**). Meis2+ neurons tagged at E12.5 are found in the LAT (inset in **C**) and in the DLP of the MED (inset in **C’**). Size bar: 200 µm

## DISCUSSION

The molecular mechanisms leading to the formation and organization of the cerebellar nuclei remain largely unknown. Here, we present a detailed fate mapping analysis covering the anatomical and molecular organization of the cerebellar nuclei in space and time during embryonic development. By GIFM, we describe the molecular trajectories linking Atoh1+ progenitors to their molecularly defined glutamatergic derivatives in the prenatal CN.

### Time window and spatial progression of glutamatergic neurogenesis

Glutamatergic CN neurons emerge as *Atoh1+* progenitors from the rostral rhombic lip and, migrating beneath the pial surface of the cerebellar anlage, reach an area called nuclear transitory zone (NTZ) by E14.5. From this location, they invade the prospective CN territory. Tamoxifen-controlled Atoh1::CreER^T2^ tracing shows neurons of the glutamatergic lineage in the putative CN starting from E9.5, although our data confirm prior findings indicating that the bulk of CN neurogenesis takes place between E10.5 and E12.5 ^17,18^. Our findings reveal that CN neurogenesis proceeds from rostral to caudal (**Fig.1A-B’**), and from lateral to medial, with the earliest *Atoh1*+ progenitors populating the prospective LAT first. Our evidence of a lateral-to-medial progression of CN neurogenesis recapitulates the findings made in the chick cerebellum, although a proper LAT is not present in that species ^1,28^.

### Brn2 is a nonspecific marker of INT and LAT neurons

In this work, we phenotyped genetically tagged progenitors with various markers known to label the cerebellar nuclei, focusing on glutamatergic neurons. One of them, namely Brn2 (Pou3F2), labels a large neuronal population almost exclusively located in the INT and LAT. However, our findings show that Brn2 also labels neurons, mostly located in ventral territories of the CN, which are never tagged by Atoh1::CreER^T2^, thus are not glutamatergic. This is confirmed by the fact that at E13.5 Brn2 colocalizes with the glutamatergic marker *Lhx9* in the NTZ, but is also found at the same stage in the cerebellar VZ, which gives rise to GABAergic neurons. This defines Brn2 as a factor affecting progenitor position from early stages of CN development, as suggested by its lateral localization in the NTZ (https://developingmouse.brain-map.org/experiment/siv?id=100058689&imageId=101146327&initImage=ish&x=3416&y=5112&z=3 ^29^), rather than a cell-fate specific marker. We speculate that Brn2 may be involved in guiding migration of VZ and RL precursors alike away from the midline and into the INT and LAT, possibly by controlling the expression of receptors sensitive to a repulsive midline cue. At any rate, many large tdT+ cells genetically tagged at E11.5 were negative for Brn2, indicating that this protein is not an exhaustive marker of INT and LAT glutamatergic neurons either (**Fig. 1F’**).

### Lmx1a and Tbr1 label partially separate glutamatergic populations in the MED

Our results indicate that glutamatergic neurons in the MED can be subdivided into Tbr1;Lmx1a double positive and Lmx1a+;Tbr1– (or Tbr1weak) subpopulations. The latter are found in the medial and dorsolateral protuberance regions of the MED. Published Cre lines tagging Lmx1a+ or Tbr1+ precursors ^30,31^ may help define the relationships between these embryonic progenitors and their adult counterparts.

### Meis2 labels a subset of glutamatergic CN neurons

Meis2 labels glutamatergic precursors throughout the CN, and become more dense in posterior territories of the CN, although many glutamatergic precursors are Meis2– in prenatal sections. Because Meis2 is a precocious marker expressed at high levels in the E11.5 subpial stream (besides the mesencephalon, https://developingmouse.brain-map.org/experiment/siv?id=100077943&imageId=101289116&initImage=ish&x=6278.358960168304&y=5708.920008626071&z=5 ^29^), likely affecting CN neuron development from early stages, we believe that the Meis2+ and Meis2– neurons observed at E18.5 belong to distinct lineages. Moreover, Meis2+ neuron distribution along the rostrocaudal and mediolateral axes of the CN seems discontinuous, possibly reflecting the organization of the cerebellar cortex in zones and stripes ^32,33^. This is in keeping with physiological data suggesting the existence of longitudinal micromodules corresponding to distinct peripheral receptive fields ^34–36^. The precise nature of adult Meis2+ vs. Meis2– glutamatergic derivatives remains unknown to date.

### Molecular heterogeneity of glutamatergic neurons

Globally, our findings point to a marked degree of heterogeneity in the glutamatergic progenitor pool at embryonic stages of CN development, in agreement with the existence of excitatory CN neuron sub-categories in the adult nuclei ^10,33,37–39^. To complete the description of this phenotypic diversity, a higher throughput and resolution will be achievable by combining traditional morphology with the application of more recent spatial transcriptomics ^40,41^ and barcoding techniques ^42^. A question that emerges from our analysis and others is whether this complexity mirrors a functional and connectomic heterogeneity, as already shown for the fastigial nucleus inter alia ^23^. The availability of Cre lines, intersectional genetics techniques ^43^ and viral constructs for stereotaxic injection e.g. ^44^, coupled with optogenetic and chemogenetic ^45,46^ approaches will elucidate the developmental trajectories linking embryonic and postnatal phenotypes, and allow neuronal subsets in the CN to be linked to their postsynaptic targets, exploring their function in vivo.

## Materials and Methods

### Animal care

All experiments described were performed in agreement with the stipulations of the San Raffaele Scientific Institute Animal Care and Use Committee (I.A.C.U.C.)

### Mouse genetics

Atoh1-CreER^T2^/0 mouse line. The Atoh1-CreER^T2^/0 mouse line was provided from the Jackson Laboratory (Tg(Atoh1-Cre/Esr1*)14Fsh/J, Stock No: 007684). These transgenic mice express a tamoxifen-inducible Cre recombinase in the endogenous promoter of Atoh1 gene, whose activity, when induced, is observed in neural progenitors of the cerebellar rhombic lip and dorsal hindbrain. ^18,47^. For genotyping: DNA was extracted from the tail of embryos or mice at P12 using Xpert directXtract Lysis Buffer Kit (Grisp Research Solutions, Biocell) and a PCR was carried out using the following primers to test the presence of Atoh1-CreER^T2^ transgene:

-oIMR1084 (Forward): 5’ GCG GTC TGG CAG TAA AAA CTA TC 3’

-oIMR1085 (Reverse): 5’ GTG AAA CAG CAT TGC TGT CAC TT 3’

The amplification product is verified through an agarose gel (1,5% in TAE 1X) electrophoresis. The expected band is 100bp for Atoh1-CreER^T2^/0.

#### Rosa26tdTomato/+ mouse line

The Rosa26tdTomato is a congenic mouse line on C57BL/6J background and was provided from the Jackson Laboratory (B6.Cg-Gt(ROSA)26sor Tm14(CAG-tdTomato)Hze/J, Stock #007914). This strain contains a loxP-flanked STOP cassette preventing the transcription of a CAG promoter-driven red fluorescent protein (RFP) variant (tdTomato) inserted into the Gt(ROSA)26Sor locus. After Cre-mediated recombination, this strain expresses robust tdTomato fluorescence. <u>Mouse lines crossing.</u> In this project, all the experiments have been performed on embryos resulting from the crossing of Atoh1-CreERT2/0 and Rosa26tdT+/+ mice. <u>Tamoxifen administration.</u> Tamoxifen (Sigma-Aldrich, T5648) has been dissolved in corn oil (Sigma-Aldrich, C8267) at a concentration of 10mg/ml by shaking overnight at 37°. Tamoxifen (approximately 80 mg/kg mouse body weight) have been delivered by oral gavage to pregnant dams at specific days post-coitum (dpc). In case of experiments on adult mice, pups were taken through a Cesarean section at E18.5 and given to an adoptive mother.

### Tissue preparation and processing

Embryos and mice were respectively fixed for 6-8 h by immersion with 4% PFA in 1× PBS, or anesthetized using Avertin (Sigma) and transcardially perfused with 0.9% NaCl followed by 4% PFA in 1× PBS. The tissues were cryoprotected overnight in 30% sucrose in 1× PBS, embedded in a solution made by 3:1 OCT (Bioptica) and 30% sucrose, and stored at −80°C. For immunofluorescence staining, 14 µm thick sagittal or rostral sections were cut using a Leica cryotome.

### Immunofluorescence staining

Sections were washed in 1× PBS to remove OCT and, if necessary, antigen retrieval was performed by heating sections in a 10mM citrate solution, pH 6, in a microwave for 5’. Sections were blocked/permeabilized in 10% goat serum, 0.3% Triton X-100 in 1× PBS for 1h and then incubated with primary antibodies overnight at 4°C. After 3 washing in 1× PBS, sections were incubated with secondary antibodies (1:500, Molecular Probes Alexa Fluor, Life Technologies) at room temperature for 2h, washed 3 times in 1× PBS, counterstained with 4′,6-diamidino-2-phenylindole (DAPI, Sigma, 1:2000 in 1× PBS) for 5’ and mounted with fluorescent mounting medium (Dako). The Lmx1a antibody detection capability was improved using the Tyramide Signal Amplification kit (TSA, Perkin Elmer), in accordance with manufacturers instructions.

### Antibodies

Brn2, rabbit, 1:500, Cell Signaling, Catalog number 12137, requires antigen unmasking; Lmx1a, rabbit, 1:3000, Millipore, Catalog number 550274, requires requires antigen unmasking and TSA; mCherry, chicken, 1:500, Novus Biologicals, Catalog number NBP2-25158; Tbr1, rabbit, 1:3000 Abcam, Catalog number Ab31940, requires antigen unmasking; Tbr1, mouse, 1:100, Catalog number 66564-1-Ig, Proteintech, requires antigen unmasking; Meis2, mouse, 1:400, Catalog number sc-515470, Santa Cruz Biotechnology.

### Microscopy and image processing

#### Fluorescence microscopy

Optical microscopy was carried out using a Mavig RS-G4 confocal microscope with 20x magnification. Quantitative and morphometric evaluations were performed with the ImageJ software (NIH), using the selection tool to delimit areas of interest. Automatic cell count was used only for Brn2+ cells: the green channel was isolated and filters Median and Watershed were applied to images to reduce background and to separate single cells, respectively. For the manual cell count, the plugin Cell Counter was used. Digital images were processed with Adobe Photoshop to adjust contrast and brightness. <u>Light microscopy</u> Light microscopy was carried out using Zeiss AxioImager M2m microscope with 2,5x magnification to analyze in situ hybridizations of bright field signals.

##### Abbreviations

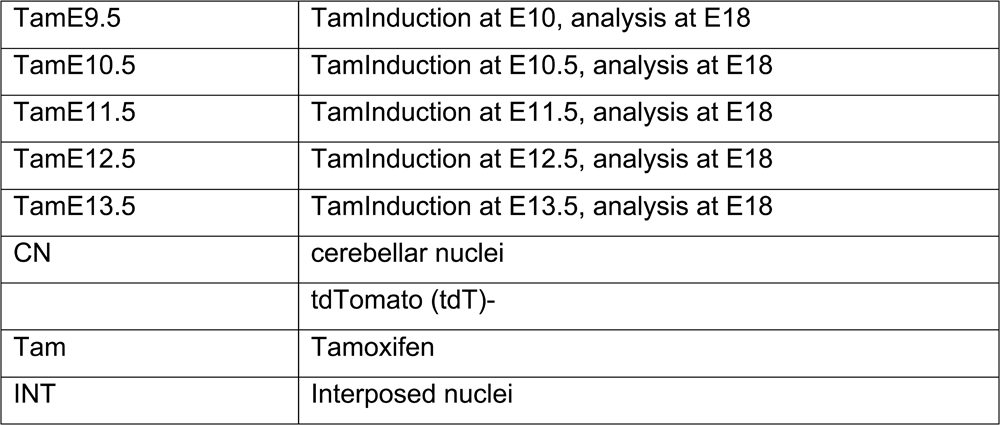
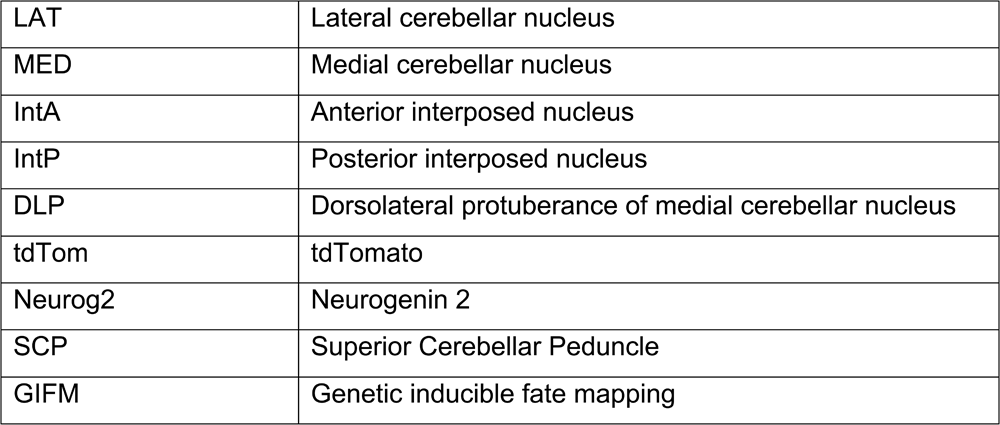

